# Three classes of response elements for human PRC2 and MLL1/2-trithorax complexes

**DOI:** 10.1101/232686

**Authors:** Junqing Du, Brian Kirk, Jia Zeng, Jianpeng Ma, Qinghua Wang

## Abstract

Polycomb group (PcG) and trithorax group (TrxG) proteins are essential for maintaining epigenetic memory in both embryonic stem cells and differentiated cells. To date, how they are localized to hundreds of specific target genes within a vertebrate genome had remained elusive. Here, by focusing on short *cis*-acting DNA elements of single functions, we discovered, for the first time, to our knowledge, three classes of response elements in human genome: PcG response elements (PREs), MLL1/2-TrxG response elements (TREs) and PcG/TrxG response elements (P/TREs). We further demonstrated that, in contrast to their proposed roles in recruiting PcG proteins to PREs, YY1 and CpG islands are specifically enriched in TREs and P/TREs, but not in PREs. The three classes of response elements as unraveled in this study open new doors for a deeper understanding of PcG and TrxG mechanisms in vertebrates.

## Introduction

Polycomb group (PcG) and trithorax group (TrxG) proteins were originally discovered in *Drosophila* where their mutations resulted improper body plans (Duncan, 1982; Ingham and Whittle, 1980; Jürgens, 1985; Kuzin et al., 1994; Lewis, 1949). Their crucial importance in development is exemplified by early lethality in mouse embryos upon deleting some of these proteins (Margueron and Reinberg, 2011). PcG proteins are present as two major multi-protein complexes, polycomb repressive complex (PRC) 1 and 2. Of them, PRC2 catalyzes the hallmark repressive mark of histone 3 Lys27 trimethylation (H3K27me3). Human PRC2 contains the catalytic subunit Enhancer of Zeste Homolog 1 or 2 (EZH1/2), Suppressor of Zeste 12 homolog (SUZ12), and Embryonic Ectoderm Development (EED) as the core subunits (Aranda et al., 2015; Bauer et al., 2016; Chittock et al., 2017; Comet et al., 2016; Geisler and Paro, 2015; Kassis and Brown, 2013; Margueron and Reinberg, 2011; Piunti and Shilatifard, 2016; Schuettengruber et al., 2017; Simon and Kingston, 2013; Whitcomb et al., 2007). TrxG proteins are also present as several multi-subunit COMPASS (complex of proteins associated with Set1) –like complexes. *Drosophila Trithorax* (*Trx*) within the COMPASS-like complex is responsible for the placement of the hallmark activation marks of H3K4me3 on genes containing PcG and TrxG response elements (PRE/TREs) (Ringrose and Paro, 2007). Mixed-lineage leukemia (MLL) 1/2 are the human homologues of *Drosophila Trx* (Geisler and Paro, 2015; Kingston and Tamkun, 2014; Piunti and Shilatifard, 2016; Schuettengruber et al., 2017).

From *Drosophila* to humans, PcG and TrxG complexes regulate the expression of hundreds of developmentally important, evolutionarily conserved target genes in each genome in a highly sequence-specific fashion. Principles for genomic targeting of these complexes initially emerged from earlier studies on PRE/TREs in *Drosophila HOX* clusters (Bauer et al., 2016; Entrevan et al., 2016; Ringrose and Paro, 2007; Schuettengruber et al., 2017; Simon and Kingston, 2013). *Drosophila* PcG and TrxG complexes were frequently found to co-occupy these PRE/TREs (Chang et al., 1995; Chinwalla et al., 1995; Gindhart and Kaufman, 1995; Orlando et al., 1998; Strutt et al., 1997), and switch from repressed to activated states were observed after a brief transcription activity (Cavalli and Paro, 1998, 1999; Maurange and Paro, 2002; Rank et al., 2002). Therefore, these elements are also termed “cellular memory modules” (CMMs). A minimal CMM of 219bp was identified in *Drosophila Fab-7* region that regulates the *Abdominal-B* gene (Dejardin and Cavalli, 2004). *Drosophila* PRE/TREs contain complex combinations of recognition motifs for transcription factors such as PHO, En1, Dsp1, Sp1, KLF, Grh, GAF and Combgap (Dejardin et al., 2005; Ray et al., 2016; Ringrose and Paro, 2007).

Despite the high conservation of PcG and TrxG complexes as well as their target genes from *Drosophila* to humans, the PRE/TREs that are signatures for their target genes are not evolutionarily conserved, which has severely halted the study of PcG and TrxG-mediated epigenetic regulation in vertebrates (Bauer et al., 2016). The first two reported vertebrate PRE/TREs were *Kr*^9kb^ and *Kr*^3kb^ in mouse genome that were found to recruit PcG in *Drosophila* and mouse cells and modulate reporter gene expression in a PcG and TrxG-dependent manner (Sing et al., 2009), and a 1.8-kb D11.12 (between human *HOXD11* and *HOXD12*) that endorsed PcG-dependent gene silencing (Woo et al., 2010).

Despite these and other pioneering studies (comprehensively reviewed in (Bauer et al., 2016)), the mechanisms of PcG and TrxG recruitment in vertebrates remained elusive and many controversies existed. For instance, some studies suggested a role of CpG islands in recruiting PRC2 complex to vertebrate PREs (Basu et al., 2014; Jermann et al., 2014; Ku et al., 2008; Lynch et al., 2012; Mendenhall et al., 2010; Tanay et al., 2007). Indeed, insertion of GC-rich and CpG-rich sequences (1,000bp) at ectopic sites was shown to be sufficient to lead to both H3K4me3 and H3K27me3 marks (Wachter et al., 2014). However, many other studies argued against such a role for CpG islands (Schorderet et al., 2013; Sing et al., 2009; van Heeringen et al., 2014). In addition, although PHO was believed to participate in recruiting PcG proteins in *Drosophila*, the roles of its mammalian homologue Yin Yang 1 (YY1) in this capacity have been under heated debate. While YY1 was frequently used as a criterion to associate with PcG function, its binding motif was found to be depleted in mammalian polycomb domains (Ku et al., 2008; Liu et al., 2010).

We reasoned that the source of the longstanding difficulties and controversies in the field is likely the large sizes of *cis*-acting DNA segments used in previous studies. We further postulated that within each *cis*-acting DNA segment, there exist short, modular building blocks of *cis*-acting DNA elements. Each such *cis*-acting DNA element performs a single function and harbors a specific signature for recognition by PcG and/or TrxG recruitment factors. Discovery and detailed characterization of these modular units will provide the foundations towards a unified mechanistic understanding of PcG and TrxG recruitment and regulation that had been so far unattainable.

In this study, by taking a “reductionist” approach, we have obtained multiple lines of solid evidence that collectively discovered, for the first time, three classes of PRE/TREs in human genome: PREs, TREs and P/TREs. In this context, PRE/TREs are used as a collective term for all response elements recognized by either PcG or TrxG and contain three classes: PREs are defined as short *cis*-acting DNA elements that are only occupied by PcG proteins (without TrxG proteins) and contain only H3K27me3 marks (without H3K4me3 marks), TREs are those that are only occupied by TrxG proteins (without PcG proteins) and contain only H3K4me3 marks (without H3K27me3 marks), and P/TREs are those that attract both PcG and TrxG complexes and harbor both H3K4me3 and H3K27me3 marks. Furthermore, this “reductionist” approach allowed unequivocal demonstrations that YY1 and CpG islands co-localize to TREs and, to a lesser extent, to P/TREs, but not PREs, thus resolving the central controversies regarding the roles of YY1 and CpG islands in recruiting human PRC2 complex.

## Results

### Prediction of candidate human PRE/TREs by *EpiPredictor*

We first used *EpiPredictor*, a bioinformatics tool that we developed earlier (Zeng et al., 2012), to predict human candidate PRE/TREs. Due to the lack of well-characterized human transcription factors known to recruit PcG or TrxG complexes, we started from *Drosophila* transcription factors with these functions, PHO, En1, Dsp1, Sp1, KLF, Grh, GAF, and gathered their known or suspected human homologues (**Table S1**). The DNA recognition motifs of these human homologues were used to scan human genome. The 61 top-ranked DNA fragments (**Table S2**) were selected for experimental investigations.

### Characterization of candidate PRE/TREs by dual luciferase reporter assay

To determine whether these top-ranked predictions by *EpiPredictor* are indeed PRE/TREs that regulate gene expression, we developed a transient dual luciferase assay in cultured human HeLa cells (**Fig.1a**). The use of HeLa cells instead of embryonic stem cells was to avoid the predomination of the repressive H3K27me3 marks. In the assay, each of the top 61 predictions flanked by two FRT sites was inserted immediately upstream of the mini-promoter that controls the expression of firefly luciferase. For negative controls, we used the empty 2FRT-YY1pLuc vector (2FRT Vector) or the vector inserted with a 200bp fragment from chr1:81132056-81132255 (Gene Desert) that was not associated with any epigenetic markers or proteins in ChIP-seq tracks on Genome Brower. PRE-*Kr*^3kb^ from mouse and PRE^d10^ from human that have been reported as PREs (Schorderet et al., 2013; Sing et al., 2009) were used as positive controls. FLPe recombinase in FLP(+) samples efficiently excised out the DNA insert between the two FRT sites (**Fig. S1a**), leading to changes in firefly luciferase activity (**Fig. S1b**). The significantly increased firefly luciferase activity of PRE-*Kr*^3kb^ and PRE^d10^ in FLP(+) samples confirmed their PRE-like behaviors and validated our transient dual luciferase assay in cultured human HeLa cells (**Fig.1b**). With this assay, we found that 19 (31.1%) DNA inserts behaved like PREs that significantly repressed the firefly luciferase reporter gene expression, and 16 (26.2%) DNA inserts that behaved like TREs to activate the expression of firefly luciferase in HeLa cells (**Fig.1b,** **Fig. S1b**). These together accounted for about 57% of the 61 top-ranked predictions.

**Fig.1.**
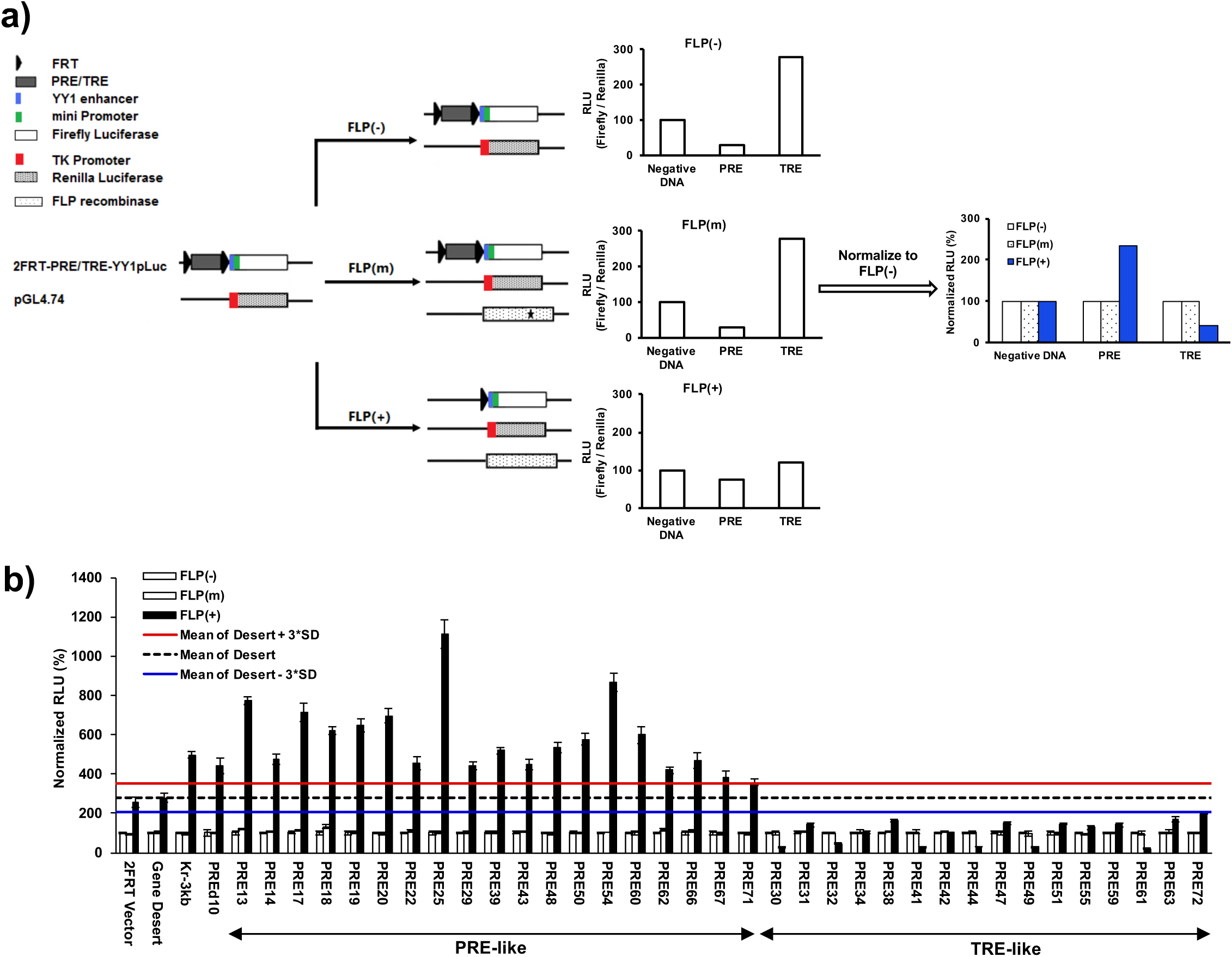
Screening of candidate PRE/TREs by dual luciferase reporter assay. **a).** Overview of the screening strategy. The assay used two plasmids: 2FRT-YY1pLuc and pGL4.74. Each of the candidate PRE/TREs was inserted between the two FRT sequences in 2FRT-YY1pLuc to yield 2FRT-PRE/TRE-YY1pLuc. pGL4.74 contains a Renilla luciferase gene under the control of an upstream thymidine kinase (TK) promoter. This allowed the signal of firefly luciferase to be normalized against that of Renilla luciferase from the same sample, yielding relative luciferase activity (RLU), to account for any variations in transfection and expression. In FLP(+) samples, an additional plasmid harboring the wild-type FLPe recombinase gene was co-transfected with the two luciferase vectors to allow the excision of candidate PRE/TRE at the two FRT sites. In FLP(m), a plasmid containing a C-terminally truncated, inactive FLPe was used in the transfection to serve as a negative control. The RLU values of the FLP(m) and FLP(+) samples were normalized against that of the FLP(−) for the same DNA insert to give rise to Normalized RLU. As PREs or TREs were supposed to repress or activate gene expression, respectively, the excision of inserted DNA fragments by FLPe in FLP(+) samples should increase the firefly luciferase signals for PREs, and decrease the firefly luciferase signals for TREs. **b).** Results of dual luciferase assay in HeLa cells. The data were represented as mean ± SD (standard deviations) from at least three replicates. The mean signal of the Gene Desert FLP(+) sample and its plus and minus 3*SD were shown as black dashed line or red or blue solid line, respectively. Candidate PRE/TREs were selected as PRE-like or TRE-like if the Normalized RLU values of their FLP(+) samples were above the red line or below the blue line, respectively. See also **Fig. S1**.

### Recruitment of PRC2 and MLL1/2-TrxG complexes to endogenous genomic loci

Next, we examined whether the DNA fragments with PRE- or TRE-like behaviors in the dual luciferase assay carry corresponding H3K27me3 or H3K4me3 marks at the endogenous loci and are enriched with key components of PRC2 or MLL1/2-TrxG complexes that catalyzes these marks. Among the 19 DNA fragments with PRE-like behavior in the dual luciferase assay, seven of them (PRE14, 17, 19, 29, 39, 48 and 62) had more than two-fold enrichment for H3K27me3 over that of Gene Desert (**Fig.2a**). In addition, all except one (PRE62) had very low enrichment for H3K4me3 marks, confirming their PRE-like behaviors (**Fig.2a**). Furthermore, among the seven PRE-like DNA fragments with strong H3K27me3 signals, five (PRE14, 29, 39, 48 and 62) had strong ChIP signals for both EZH2 and EED (**Fig.2b**). Indeed, in HeLa cells in which the expression of SUZ12 was knocked down using shRNA lentiviruses (**Fig.2c**), all PRE14, 29, 39, 48 and 62 as well as PRE-*Kr*^3kb^ exhibited statistically significantly higher luciferase activity than those in cells with scrambled shRNA (**Fig.2d**). In addition, the endogenous loci of PRE14, 29, 39, 48 and 62 all showed statistically significant decrease of EED signals upon SUZ12 knockdown (**Fig.2e**).

**Fig.2.**
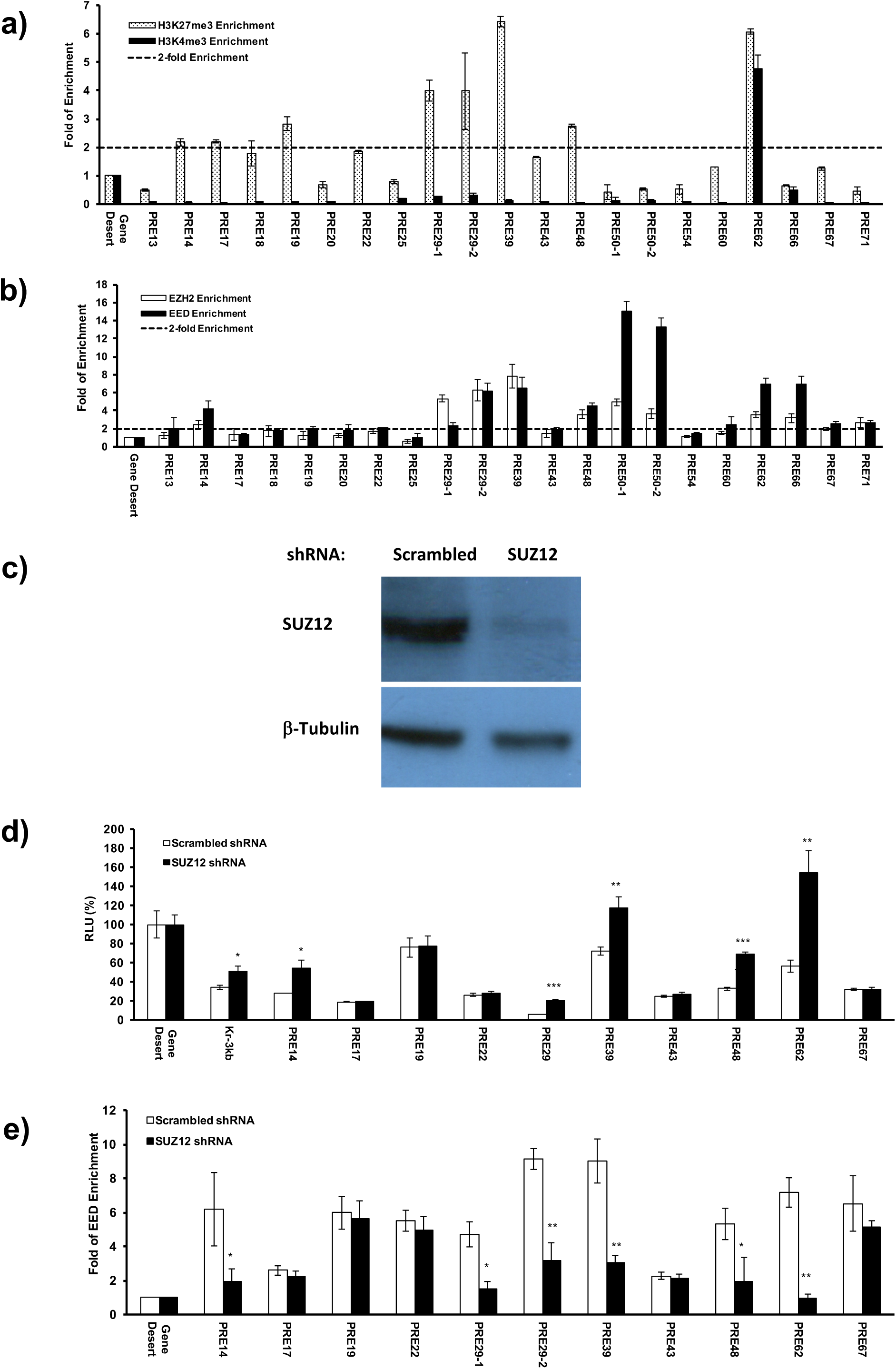
Enrichment of H3 marks and PRC2 at the endogenous loci of PRE-like DNA fragments. **a).** The enrichment of H3K27me3 (dotted open bars) and H3K4me3 (filled solid bars) marks at the endogenous loci of PRE-like DNA fragments in HeLa cells assessed by ChIP-qPCR. **b).** The enrichment of EZH2 (open bars) and EED (filled solid bars) measured by ChIP-qPCR. **c).** The expression of SUZ12 in SUZ12 shRNA knockdown cell line was significantly reduced as compared to that in control cell line treated with scrambled shRNA. **d).** The luciferase activity of PRE-like DNA fragments in SUZ12 knockdown cells (filled solid bars) compared to control cell line treated with scrambled shRNA (open bars). The RLU at Gene Desert was considered as 100%. **e).** The enrichment of EED at the endogenous loci of PRE-like DNA fragments in SUZ12 knockdown cell line. In **a, b, e,** the enrichment at Gene Desert was treated as 1.0. In **a, b, d, e,** data were represented as mean ± SD from at least three replicates. The *p* values of Student’s *t*-test comparing SUZ12 shRNA and scrambled shRNA (**d, e**) were showed as * for *p*<0.05, ** for *p*<0.005 and *** for *p*<0.001.

On the other hand, of the 16 DNA fragments with TRE-like behavior in the dual luciferase assay, all had low signals for H3K27me3 (**Fig.3a**), confirming their TRE-like characteristics. Nine of them (PRE30, 32, 34, 41, 42, 44, 49, 55, and 61) had at least two-fold enrichments for H3K4me3 compared to that of Gene Desert (**Fig.3a**). Among them, eight DNA fragments (PRE30, 34, 41, 42, 44, 49, 55 and 61) had at least 2-fold enrichment in ChIP signals for both MLL1 and WD repeat-containing protein 5 (WDR5), which is a key component of human COMPASS complexes (**Fig.3b**). Additionally, in WDR5-knockdown HeLa cells (**Fig.3c**), seven TRE-like DNA fragments (PRE30, 34, 41, 42, 44, 55 and 61) exhibited significant decrease in luciferase activity (**Fig.3d**), and six of them (PRE30, 41, 42, 44, 55 and 61) showed significantly reduced enrichment of MLL1 at their endogenous loci than in cells with scrambled shRNA (**Fig.3e**).

**Fig.3.**
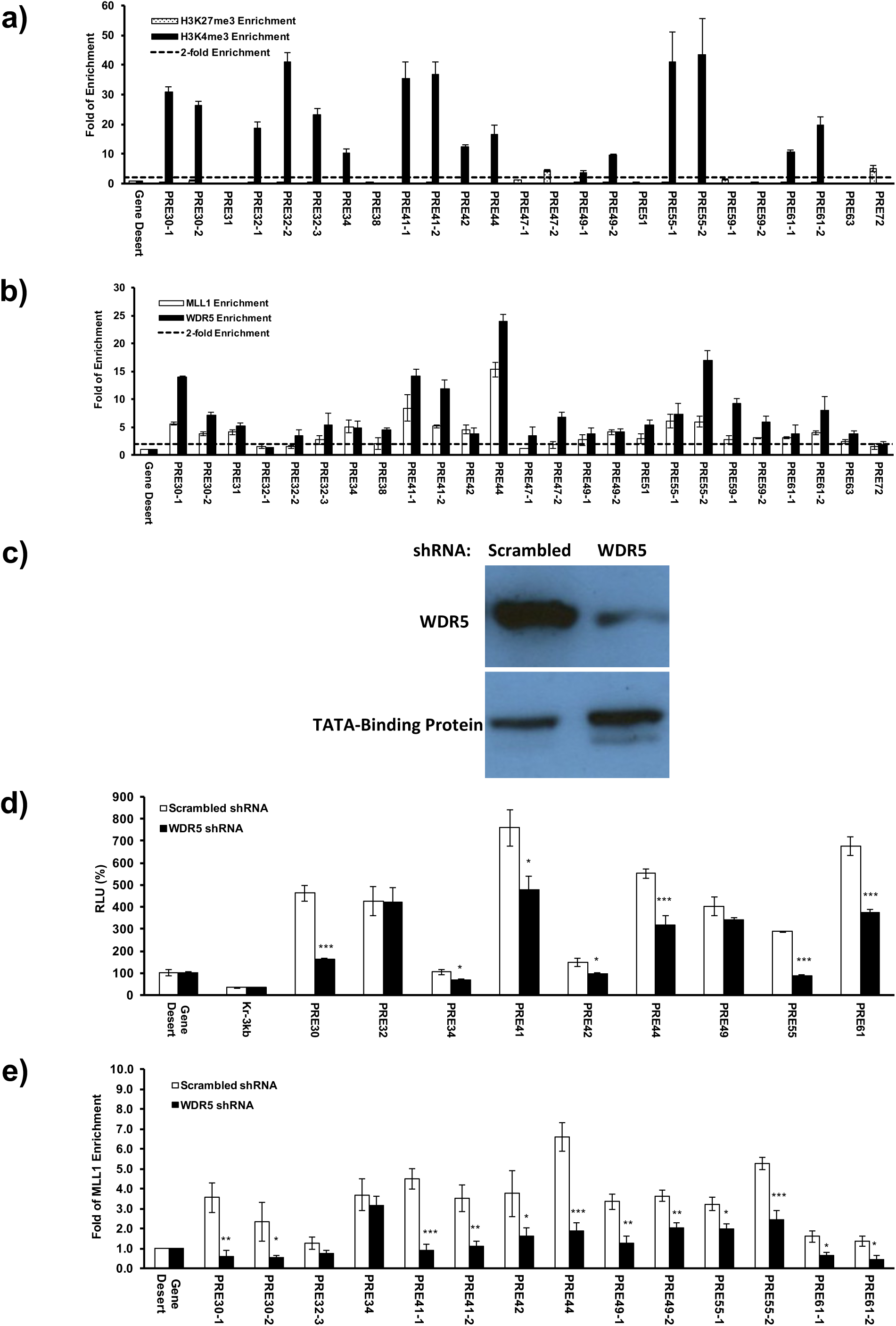
Enrichment of H3 marks and MLL1/2-TrxG complex at the endogenous loci of TRE-like DNA fragments. **a).** The enrichment of H3K27me3 (dotted open bars) and H3K4me3 (filled solid bars) marks at the endogenous loci of TRE-like DNA fragments in HeLa cells assessed by ChIP-qPCR. **b).** The enrichment of MLL1 (open bars) and WDR5 (filled solid bars) measured by ChIP-qPCR. **c).** The expression of WDR5 in WDR5 shRNA knockdown cell nuclear extract was significantly reduced as compared to that in control cell line treated with scrambled shRNA. **d).** The luciferase activity of TRE-like DNA fragments in WDR5 knockdown cells (filled solid bars) compared to control cell line treated with scrambled shRNA (open bars). The RLU at Gene Desert was considered as 100%. **e).** The enrichment of MLL1 at the endogenous loci of TRE-like DNA fragments in WDR5 knockdown cell line. In **a, b, e,** the enrichment at Gene Desert was considered as 1.0. In **a, b, d, e,** data were represented as mean ± SD from at least three replicates. The *p* values of Student’s *t*-test comparing WDR5 shRNA and scrambled shRNA (**d, e**) were showed as * for *p*<0.05, ** for *p*<0.005 and *** for *p*<0.001.

Collectively, we have identified five putative PREs (PRE14, 29, 39, 48 and 62) that were enriched for PRC2 complex, had a high H3K27me3 level and repressed luciferase expression, and six putative TREs (PRE30, 41, 42, 44, 55 and 61) that were enriched for MLL1/2-TrxG complex, had a high H3K4me3 level and activated luciferase expression. However, PRE62 was different from all other putative PREs: it also had a high level of H3K4me3 enrichment at the endogenous locus.

### Identification of core sequences of putative PREs and TREs

We selected five putative PREs (PRE14, 29, 39, 48 and 62) and four putative TREs (PRE30, 41, 44, 55) for further investigation. These DNA fragments were of the size of 360-700bp long. Previous studies have shown that mouse DNA sequences as short as 220bp can recruit H3K27me3 marks (Jermann et al., 2014). We asked whether these DNA fragments can be further truncated while maintaining the observed PRE or TRE properties. We tested the luciferase activity of multiple truncations and identified the functional core for each individual DNA fragment (**Fig.4a**). The core DNA sequence was defined as the shortest fragment with the highest activity in repression (for putative PREs) or activation (for putative TREs) among all tested truncations for a given DNA fragment. Sometimes, the core DNA sequences even had higher activity than the full-length DNA fragment, examples including PRE14 and PRE44 (**Fig.4a**). These core DNA sequences were in the range of 113-266bp for putative PREs and 170-348bp for putative TREs. Therefore, the repression or activation activities of these putative PREs or TREs, respectively, were localized in short DNA fragments.

**Fig.4.**
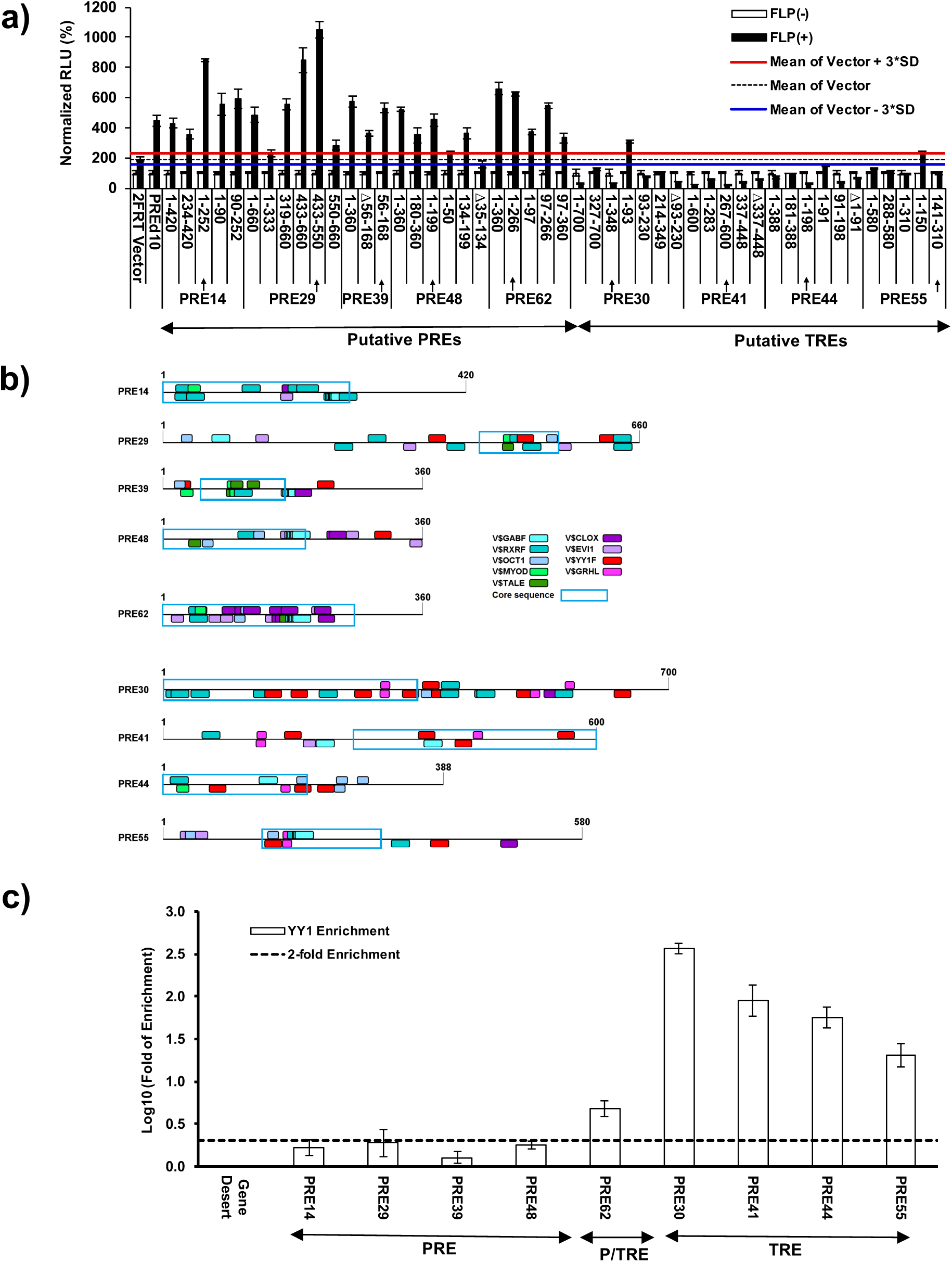
Define the core sequences of putative PREs and TREs and transcription factor binding motifs. **a).** Normalized RLU of various truncations of putative PREs and TREs. The arrows indicated the regions corresponding to the core sequences. For each sample, the signals of FLP(−) was deemed as 100% against which the signal of the FLP(+) sample was normalized. The data were represented as mean ± SD from at least three replicates. The mean signal of the 2FRT Vector FLP(+) sample and its plus and minus 3*SD were shown as black dashed line or red or blue solid line, respectively. The firefly luciferase signals were considered as significantly changed if the Normalized RLU values were above the red line or below the blue line. **b).** Distribution of transcription factor binding motifs in the three classes of PRE/TREs. **c).** Enrichments of transcription factor YY1 at the endogenous loci of the nine PRE/TREs where the enrichment at Gene Desert is 1.0. Also see **Table S6**.

### Validation of putative PREs and TREs in identical genomic environment

PREs are sensitive to genomic location (Okulski et al., 2011; Steffen and Ringrose, 2014). To further validate these putative PREs and TREs in a well-controlled genomic environment, we integrated each core sequence with luciferase reporter cassette into the AAVS1 site (the adeno-associated virus integration site 1) in human chromosome 2 by the clustered, regularly interspaced, short palindromic repeats and Cas9 endonuclease (CRISPR-Cas9) system (**Fig.5a,** **Fig. S2**). AAVS1 has an open chromatin structure, is transcription-competent and represents a well-validated genomic location for testing cellular functions of a DNA fragment (Sadelain et al., 2011). Most importantly, there is no known adverse effect on the cells from the DNA fragment inserted at AAVS1. Following Cas9 cleavage and homology-directed repair (HDR), the colonies with desired inserts were selected by flanking PCR and verified by Sanger sequencing (**Fig. S2)**. Dual luciferase activities were measured for a representative clone of each core sequence (**Fig.5b**). Clearly, the FLP(+) samples of cells stably carrying PRE14, 29, 39, 48, 62 core sequences at the AAVS1 site exhibited significantly higher expression of firefly luciferase while those of cells integrated with PRE30, 41, 44, 55 core sequences showed significantly reduced luciferase expression than their FLP(-) samples (**Fig.5b**). Furthermore, except for PRE62, all putative PREs (PRE14, 29, 39 and 48) had significant enrichment for EZH2 and H3K27me3 and no enrichment for WDR5 or H3K4me3. On the contrary, all putative TREs (PRE30, 41, 44 and 55) had no enrichment for EZH2 or H3K27me3 but significant enrichment for WDR5 and H3K4me3 (**Fig.5c**). Strikingly, PRE62 again exhibited a unique behavior for being enriched for all EZH2, H3K27me3, WDR5 and H3K4me3. These data provided additional and final proof that we have discovered three classes of PRE/TREs in human HeLa cells: PREs such as PRE14, 29, 39 and 48, TREs such as PRE30, 41, 44 and 55 and P/TRE such as PRE62. In addition, we also included PRE^d10^ as a positive control for its shorter sequence (1.4kb) than PRE-*Kr*^3kb^. Clearly, PRE^d10^ behaved like PRE62 with significant enrichment for all EZH2, H3K27me3, WDR5 and H3K4me3, and exhibited repressed luciferase activity (**Fig.5b, c**). Future studies are needed to investigate whether the bivalent phenotype of PRE^d10^ is localized to a shorter DNA region.

**Fig.5.**
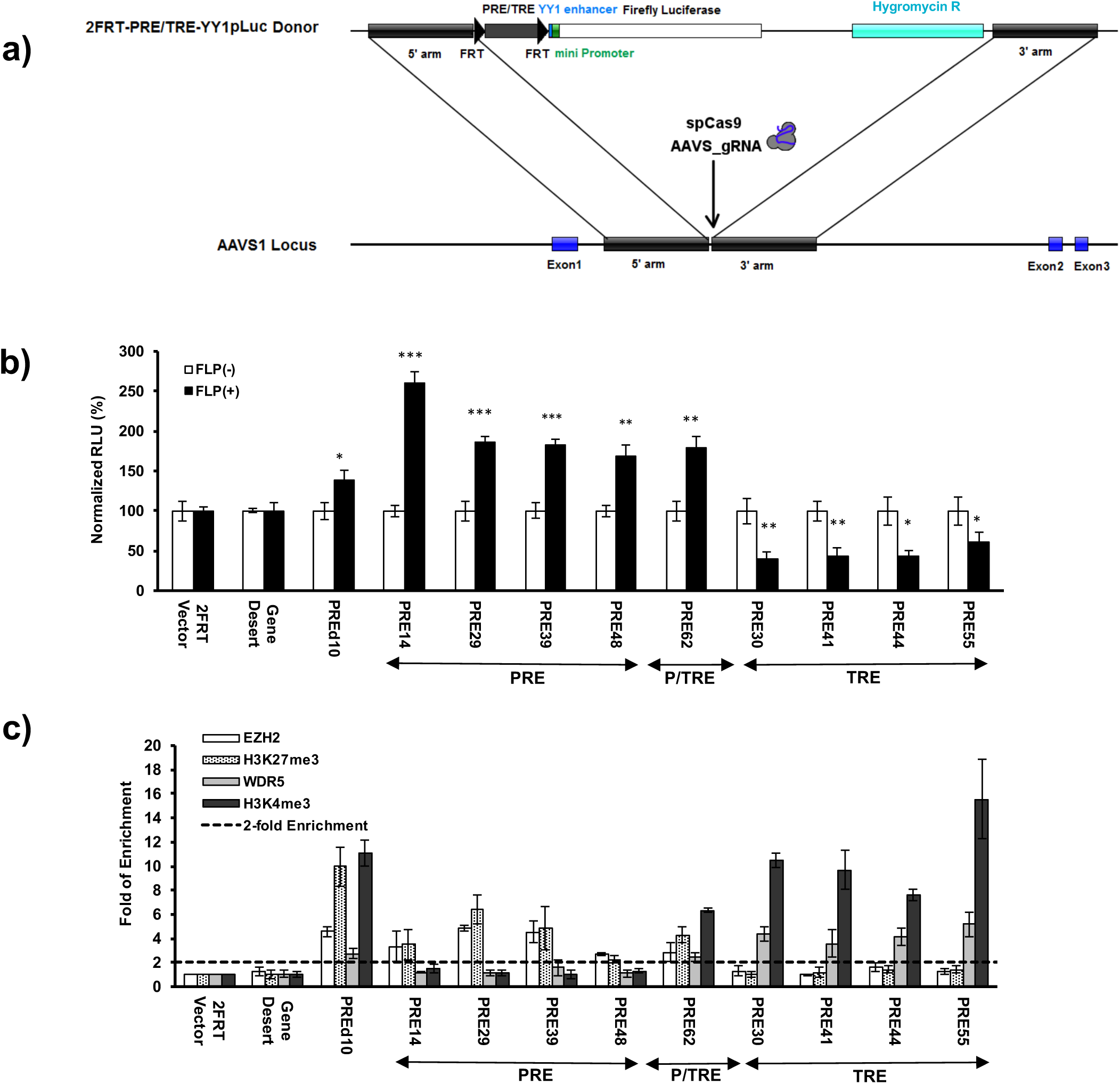
Effects of putative PREs and TREs on transcription in an identical genomic environment. **a).** Schematic illustration of the strategy that integrated the core sequences of the putative PREs and TREs (2FRT-PRE/TRE-YY1pLuc donor) into AAVS1 locus by CRISPR-Cas9. **b).** Dual luciferases assay in HeLa cells carrying stably integrated core sequences. For each sample, the signals of FLP(−) was deemed as 100% against which the signal of the FLP(+) sample was normalized to yield Normalized RLU. Data are represented as mean ± SD from at least three replicates. The *p* value of Student’s *t*-test comparing FLP(+) and FLP(−) for each integrated core sequence was shown as * for *p*<0.05, ** for *p*<0.005 and *** for *p*<0.001. **c).** The enrichment of histone marks and EZH2 and WDR5 at the promoter region downstream of the integrated core sequences. The 2-fold enrichment was shown as a dashed line. Also see **Fig. S2**.

### Identification of endogenous target genes

The next questions we asked were whether each class of response elements has specific features and what their target genes are? Toward this end, we first uncovered that the GC contents of these core sequences were significantly distinct from each other. In particular, the four PREs (PRE14, 29, 39 and 48) had an average GC content of 54.4±2.8%, the single P/TRE (PRE62) had a GC content of 47%, while the four TREs (PRE30, 41, 44 and 55) had an average GC content of 66.0±4.2% (with a *p* value of 0.0035 from the four PREs in two-tailed Student’s *t*-test) (**Table S4**). Furthermore, all TREs (PRE30, 41, 44 and 55) are located within or partially overlap with CpG islands, while the four PREs and the single P/TRE do not overlap with any CpG islands.

We examined the expression level of genes within ±500kb of each of these response elements by using HeLa cells harboring SUZ12 or WDR5 shRNA to discover their endogenous targets. Statistically significant increase of transcription upon SUZ12 knockdown allowed us to define the following target genes: AK023819 and LINC01819 (PRE14), carbohydrate sulfotransferase 2 (CHST2) and AK097380 (PRE29), LOC101928441 (PRE39), potassium calcium-activated channel subfamily M alpha 1 (KCNMA1) (PRE48), AK093368, AK127336 and BC127870 (PRE62) (**Fig.6a,** **Table S5**). Except CHST2 and KCNMA1, all other targets are currently annotated as long noncoding RNAs. In sharp contrast, the expression level of all genes proximal to the TRE class (PRE30, 41, 44 and 55) was unaffected by SUZ12 knockdown. On the other hand, WDR5 knockdown led to no significant expression changes for the targets of the PRE class, but significantly decreased transcription levels for the following target genes of the P/TRE and TRE classes: AK093368 and BC127780 (PRE62), AKT1 Substrate 1 (AKT1S1) and TBC1 domain family member 17 (TBC1D17) (PRE30), the nuclear transcription cofactor host cell factor 1 (HCF1) and transmembrane protein 187 (TMEM187) (PRE41), receptor of activated protein C kinase 1 (GNB2L1) (PRE44), heterogeneous nuclear ribonucleoproteins A2/B1 (HNRNPA2B1) and chromobox protein homolog 3 (CBX3) (PRE55) (**Fig.6b,** **Table S5**). All of the targets of the TRE class (PRE30, 41, 44 and 55) are housekeeping genes.

**Fig.6.**
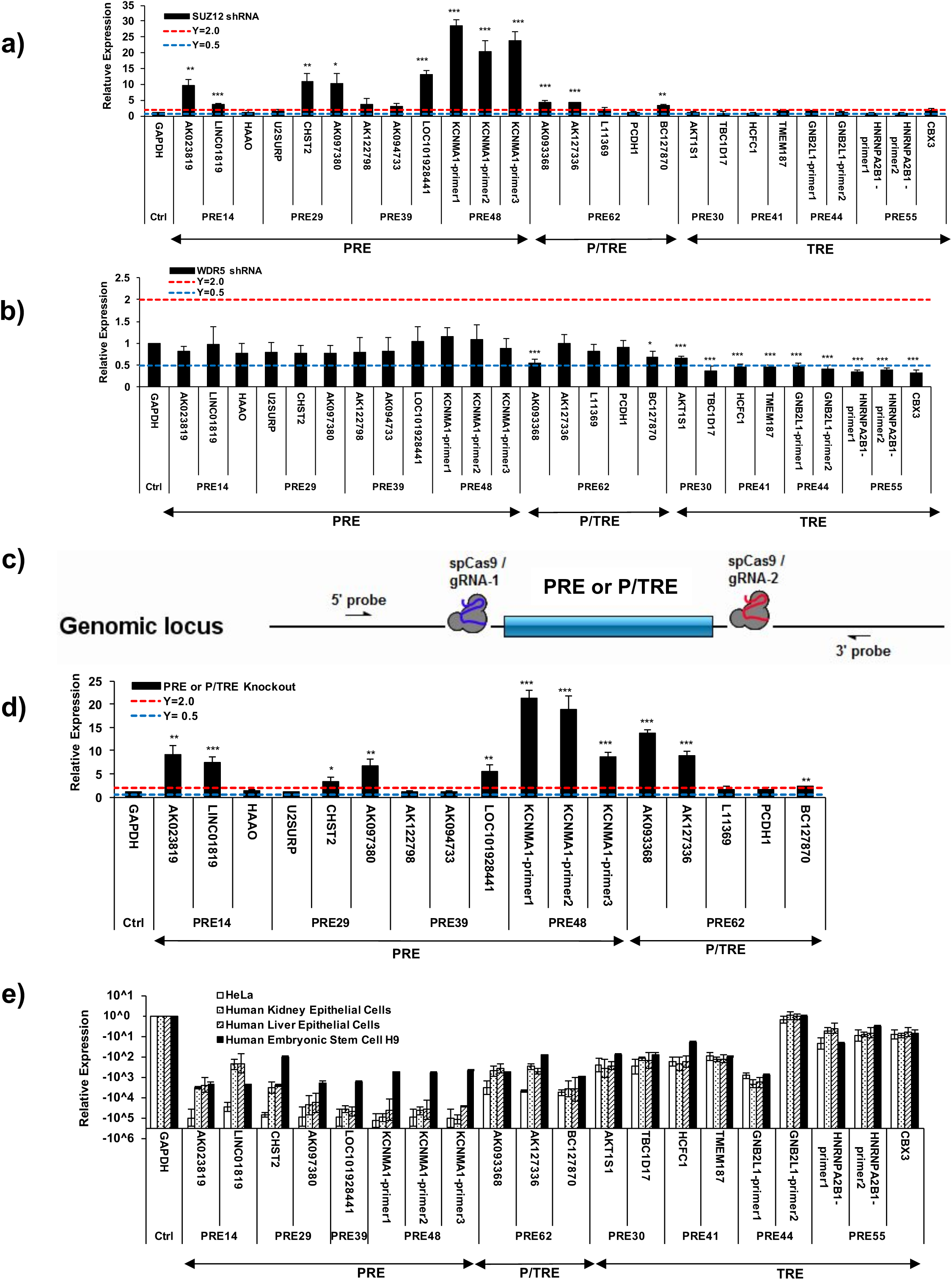
Identification of endogenous targets regulated by the nine human PRE/TREs. **a).** RT-qPCR of neighboring genes in SUZ12 knockdown HeLa cells. The genes with significantly increased transcription than in control cell line treated with scrambled shRNA were deemed as PRE or P/TRE targets. **b).** RT-qPCR of neighboring genes in WDR5 knockdown HeLa cells. The genes with significantly decreased transcription than in control cell line treated with scrambled shRNA were deemed as P/TRE or TRE targets. **c).** Schematic illustration of the strategy used to knockout PRE and P/TRE fragments from the genome loci in HeLa cells by CRISPR-Cas9. **d).** RT-qPCR of neighboring genes in PRE or P/TRE-knockout HeLa cells. **e).** RT-qPCR of neighboring genes from different cells. In all panels, the expression data were represented relative to that of housekeeping GAPDH deemed as 1.0. The *p* values of Student’s *t*-test comparing the expression levels of target genes in knockdown (**a, b**) or knockout (**d**) HeLa cells and control cells were showed as * for *p*<0.05, ** for *p*<0.005 and *** for *p*<0.001. Also see **Table S5**.

From **Table S5**, we noticed a distinct feature in terms of the distance of these three classes of PRE/TREs to the transcription start site (TSS) of the targets. While the TREs (PRE30, 41, 44 and 55) partially overlap with or are very close to the TSS of target genes, this distance is in the range of 6kb-142kb for PREs (PRE14, 29, 39 and 48) and 3-19kb for P/TRE (PRE62). The longer distance of PREs and P/TRE to TSSs allowed us to knock out the core sequences using the CRISPR-Cas9 system (**Fig.6c**). Relative to the wild-type cells, the knockout resulted in an increased transcription level of all the targets of the PRE and P/TRE classes (**Fig.6d**). These results thus provided additional evidence for the endogenous gene targets of the PREs and P/TRE classes.

Further investigation in the UCSC Genome Brower revealed that, while each of the targets of the TRE class maintains essentially a constant transcription level in cell lines of different developmental stages and tissues, the transcription levels of the targets in the PRE class change wildly. To illustrate this point, we compared the expression level of all the target genes by RT-qPCR in four cell lines, wild-type HeLa cells, human primary kidney epithelial cells, human primary liver epithelial cells, and human embryonic stem cell line H9 (**Fig.6e**). By using the expression of GAPDH in each cell line as a reference, the expression of the target genes of the PRE class (PRE14, 29, 39 and 48) were repressed to a varying degree (between 10^−2^ to 10^−5^) in different cell lines (**Fig.6e**). In sharp contrast, the target genes of PRE62 (P/TRE class) varied by ~10 folds in different cell lines, while the targets of the TRE class (PRE30, 41, 44 and 55) remained fairly constant in all cell lines (**Fig.6e**).

### Enrichment of transcription factor binding sites in the core PRE/TREs

DNA sequence-specific transcription factors are expected to be at least partially responsible for recruiting PcG and/or TrxG proteins to their genomic targets (Klose et al., 2013). However, until now we knew very little of transcription factors that perform these functions in mammalian cells. Here we used the core sequences of identified PREs, P/TRE and TREs to discover enrichment of sequence-specific transcription factors. The reason for using the core sequences of these PRE/TREs was to ensure high signal-to-noise ratio in the analysis. The analysis discovered motifs that were ubiquitously enriched in all three classes of response elements (such as the binding motif for the GABF family) (**Table S6,** **Fig.4b**). In addition, motifs recognized by families of RXRF, OCT1, MYOD, or TALE were enriched in the PRE class, those recognized by the families of CLOX and EVI1 were enriched in the P/TRE class, while those bound by the YY1F and GRHL families were uniquely enriched in the TRE class (**Table S6,** **Fig.4b**). We selected YY1, which exhibits a good level of expression in HeLa cells for experimental verification. The ChIP-qPCR results clearly demonstrated the predominant preference of YY1 at all four TREs, to a lesser degree at the P/TRE (PRE62), but not at any of the four PREs (**Fig.4c**). These data confirmed previous studies that YY1 is unlikely a recruiter of mammalian PcG complexes (Kahn et al., 2014; Mendenhall et al., 2010; Vella et al., 2012).

### Genome-wide identification and characterization of PRE/TREs in human K562 cells

The nine *cis*-regulatory DNA sequences characterized in this study have two outstanding features: **a).** they are of short sizes (113-348bp); **b).** each is enriched for histone marks (H3K27me3 for PRE, H3K4me3 for TRE, and both H3K27me3/H3K4me3 for P/TRE) and the corresponding histone modifying protein complexes. We next employed these two features to identify PRE/TREs on a genome-wide scale in human K562 cells using publically available data. By applying very stringent criteria to ensure high signal-to-noise ratio, we retrieved 15,173 PREs (416bp or smaller) that were uniquely enriched for both EZH2 and H3K27me3, 10,693 TREs (416bp or smaller) that were specifically enriched for MLL2 and H3K4me3, and 107 P/TREs that contained fully overlapping EZH2/H3K27me3 and MLL2/H3K4me3 marks (**Table S7**). The requirement of completely overlapping EZH2/H3K27me3 and MLL2/H3K4me3 marks in the 107 P/TREs was to ensure a genuine bivalent state, which was obviously a substantial underestimate of the P/TRE class in the genome. These genome-wide response elements were then used to overlap with 1,309 YY1 peaks (**Fig.7a~c**). Strikingly, in marked contrast to the high percentage of TREs that had overlapping YY1 signals, at 3.1%, only 0.02% PREs had overlapping YY1 signals (**Fig.7a,** **7b**). In addition, one of the 107 P/TREs (at 0.9%) also had overlapping YY1 signals (**Fig.7c**). Therefore, despite the small number of YY1 peaks in the original ChIP-seq data, the trend of YY1 to prefer TREs and P/TREs was very clear. Collectively, our data confirmed that, different from the roles of *Drosophila* PHO to recruit PcG to PREs, YY1 in human has specific roles in bridging the MLL1/2-TrxG complex with TREs and P/TREs.

**Fig.7.**
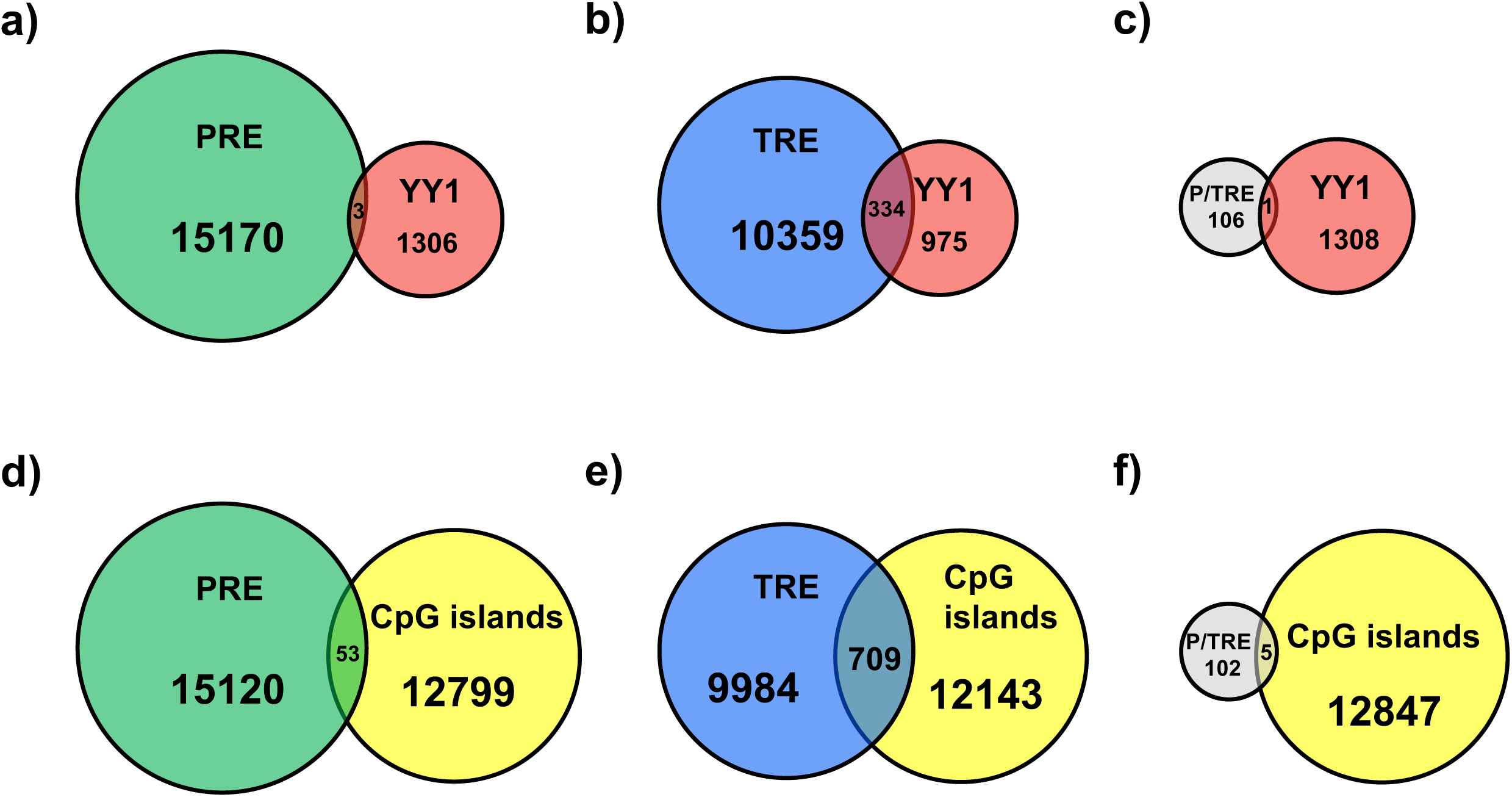
Genome-wide analysis of K562 cell line. **a~c).** Venn diagram showing the overlaps of YY1 with genome-wide PREs (**a**), TREs (**b**) and P/TREs (**c**). **d~f).** Venn diagram showing the overlaps of CpG islands with genome-wide PREs (**d**), TREs (**e**) and P/TREs (**f**). Also see **Table S7**.

We also investigated the overlap of these three classes of genome-wide PRE/TREs with 12,852 CpG islands (500bp or smaller). In marked contrast to the large percentage of TREs and P/TREs, at 6.6% and 4.7%, that overlapped with CpG islands, there was only 0.3% PREs overlapping with CpG islands (**Fig.7d~f**). These data, together with our observation on the four PREs (PRE14, 29, 39 and 48), suggest that CpG islands are unlikely key features responsible for recruiting PcG proteins to human PREs.

## Discussion

### Three classes of PRE/TREs in human genome

The three classes discovered in this study are fundamentally distinct from each other in many aspects: 1). The distance to TSS. All members of the PRE class are located at 6-142kb away from the TSS of the endogenous targets. The only P/TRE (PRE62) is 3-19kb away from the TSS of its targets, while the members of the TRE class are very close to, and often overlap with, the TSS site of their targets. 2). The GC contents. The GC contents of the four PREs (PRE14, 29, 39 and 48) were ~12% lower than those of the four TREs (PRE30, 41, 44 and 55). Similarly, we compared 105 randomly selected PREs and TREs retrieved from genome-wide analysis of human K562 cell line and found that the GC contents of the PREs were ~15% lower than those of TREs. 3). Co-localization with CpG islands. All TREs (PRE30, 41, 44 and 55) are located within or partially overlap with CpG islands, while all PREs and the single P/TRE do not overlap with any CpG islands. Similarly, in the genome-wide analysis, CpG islands were found to be specifically enriched in TREs, slightly less enriched in P/TREs, but at a very low level in PREs. 4). Their endogenous targets and expression levels. The target genes of the four PREs (PRE14, 29, 39 and 48) are predominantly long non-coding RNAs and the expression levels of which varied wildly in different cells. The target genes of the P/TRE (PRE62) are also non-coding RNAs, but their expression levels only changed within a small range in different cells. In marked contrast, the target genes of the four TREs (PRE30, 41, 44 and 55) are exclusively housekeeping genes that exhibited almost invariable expression level across different cells. 5). Recognition by transcription factors. All TREs (PRE30, 41, 44 and 55) and the single P/TRE (PRE62) were specifically enriched for YY1, but not the PREs (PRE14, 29, 39 and 48). This was also confirmed in genome-wide analysis where YY1 was found to co-localize with TREs, to a lesser degree with P/TREs, but at a very low level with PREs. All these intrinsic differences among the three classes of PRE/TREs provide strong support that they indeed carry out distinct functions and represent the basic building blocks for PcG and TrxG functions in the cell.

Whether the same three classes of PRE/TREs exist in *Drosophila* and other species awaits to be investigated. The previously identified *Drosophila* PRE/TREs in HOX clusters, also known as CMMs that are co-occupied by PcG and TrxG proteins and can switch from repressed states to activated states, behave like the P/TRE class in the three-class system of human genome. The bivalent domains found in mammalian pluripotent or multipotent cells (Azuara et al., 2006; Bernstein et al., 2006; Ku et al., 2008; Mikkelsen et al., 2007; Pan et al., 2007; Roh et al., 2006; Sanz et al., 2008; Vastenhouw and Schier, 2012; Zhao et al., 2007) likely harbor many modular P/TREs. However, the discovery of the P/TRE class in HeLa cells, together with what were reported by Maini *et al*. and Erceg *et al*. (Bengani et al., 2013; Erceg et al., 2017; Jaensch et al., 2017; Maini et al., 2017), suggests that the bivalent marks of H3K27me3 and H3K4me3 have a more general role in regulating gene expressions in embryonic stem cells, throughout the development and in differentiated cells. In addition, the TRE class of human response elements was previously unknown. Furthermore, many previously reported vertebrate PREs (reviewed in (Bauer et al., 2016)) were only probed for the enrichment of PcG proteins and H3K27me3 marks. These PREs may need to be revisited to properly classify them using the three-class system as revealed here. Thus, the discovery of the three classes of human response elements not only suggests much broader cellular functions that are regulated by PcG and TrxG proteins than previously appreciated, but also provides a new standard for characterizing vertebrate response elements.

### The role of YY1 in recruiting MLL1/2-TrxG proteins

An important unresolved issue in the PcG and TrxG field is how these protein complexes are recruited to their specific genomic targets. One central controversy was on YY1. Although its *Drosophila* homologue, PHO, has been implicated in PcG recruitment (Fritsch et al., 1999), YY1’s binding motif was puzzlingly depleted in mammalian polycomb domains (Ku et al., 2008; Liu et al., 2010). Our results from both detailed characterization of the nine PRE/TREs and genome-wide analysis of human K562 cell line clearly demonstrated that YY1 is specifically enriched at TREs and P/TREs, but not at PREs. These new findings, together with previous observations that YY1 was found to bind at active promoters in human genome but not PcG target genes (Kahn et al., 2014), suggest that, despite the extreme conservation of the DNA binding domains between YY1 and PHO, YY1 apparently has evolved a role in recruiting MLL1/2-TrxG complex, which is distinct from that of PHO in *Drosophila*.

### The role of CpG islands in recruiting MLL1/2-trithorax complexes

Another pivotal controversy in the field was the role of CpG islands in recruiting PcG complexes. Previous studies have suggested a role of CpG islands in recruiting PRC2 (Jermann et al., 2014; Ku et al., 2008; Lynch et al., 2012; Mendenhall et al., 2010;Tanay et al., 2007), while other studies argued against such a role (Schorderet et al., 2013; Sing et al., 2009; van Heeringen et al., 2014). However, all these studies examined relatively large chromatin regions. Our studies on much smaller DNA fragments (113-348bp) revealed that all four PREs (PRE14, 29, 39 and 48) do not overlap with CpG islands. In marked contrast, all TREs (PRE30, 41, 44 and 55) are located within or partially overlap with CpG islands. Furthermore, genome-wide analysis of K562 cells found that the TREs and P/TREs harbor more than 20-fold enrichment of CpG islands than the PREs. These data collectively demonstrated that CpG islands are likely involved in recruiting MLL1/2-trithorax complexes in human TRE and P/TRE sites. This conclusion agrees very well with the overwhelming genome-wide overlaps between CpG islands and H3K4me3 peaks (Orlando et al., 2012), but of minimal overlaps with H3K27me3 (Thomson et al., 2010). Previous studies suggested that insertion of GC-rich and CpG-rich sequences (1,000bp) at ectopic sites was sufficient to recruit both H3K4me3 and H3K27me3 marks (Wachter et al., 2014) and short DNA sequences (500-1,000bp) capable of recruiting H3K27me3 marks also have enrichment for H3K4me3 (Jermann et al., 2014). These observations likely reflect the cases of P/TREs where the recruitment of MLL1/2-trithorax complexes to P/TREs *via* the CpG islands somehow leads to the recruitment of PRC2 complex and the subsequent H3K27me3. However, the detailed mechanisms underlying this process await future elucidation.

### The “reductionist” approach for dissecting complex gene regulations

All the information dictating the development of a human zygote into a well-developed body is stored in the same genome that is shared by all types of cells in the body. Therefore, in order to comprehensively and precisely instruct specific gene expression in different cells, human genes carry long *cis*-regulatory DNA regions including promoters and enhancers with highly complexed organization of binding sites for transcription factors and other regulatory factors. Consequently, the use of long *cis*-regulatory DNA regions mingled with multiple functions was partially responsible for the difficulties and confusions in the field of PcG and TrxG-mediated regulation. By focusing on shorter DNA fragments of single functions in this study, the “reductionist” approach has demonstrated enormous power in distinguishing classes of fundamentally different PRE/TREs in human genome, and in unraveling the underlying DNA signatures and transcription factors required for recruitment. Future investigations into these basic building blocks of gene regulations will provide the much-needed foundations for reconciling many controversies in the PcG and TrxG field and to elucidate how various combinations of such building blocks collectively endorse robust responses in the cell.

## Methods

### Prediction of human PRE/TREs by *EpiPredictor*

An *in-silico* approach called *EpiPredictor*, which was originally trained and tested in the genome of *Drosophia* (Zeng et al., 2012), was used to predict the candidate PRE/TREs in human genome. Seven transcription factors that are known to play a role in PcG/TrxG recruitment in *Drosophia* were selected (**Table S1**) and the DNA binding motifs of their possible human homologues were used. In the prediction, we applied a sliding window to scan the entire human genome (version GRCh37/hg19) and ranked candidate PRE/TREs in a descending order based upon the scores from the support vector machine-based classifier (**Table S2**). The top 61 highly ranked predictions (PRE13-14, 16-74) were subjected to dual luciferase assay (PRE15 failed to be cloned).

### Dual luciferase reporter gene assay

Each of the predicted PRE/TRE flanked by two FRT sites was inserted into the YY1pLuc vector (Woo et al., 2010), yielding a recombinant plasmid 2FRT-PRE/TRE-YY1pLuc. The FLP(+) samples of the Gene Desert was regarded as the background for this dual luciferase assay setup for selection of DNA fragments with PRE-like or TRE-like behaviors. See more details in **Supplementary Methods**.

### Expression analysis by RT-qPCR

Total RNA was prepared using RNeasy kit (Qiagen) and reverse-transcribed using SuperScript III First Strand Synthesis kit (Invitrogen). The cDNA was used as template in qPCR. The relative expression level was compared to that of GAPDH in each cell line.

## Acknowledgements

J.M. thanks support from the National Institutes of Health (R01-GM067801, R01-GM116280), and the Welch Foundation (Q-1512). Q.W. acknowledges support from the National Institutes of Health (R01-AI067839, R01-GM116280), and the Welch Foundation (Q-1826). B.D.K and J.Z. were partially supported for postdoctoral training fellowships from the Keck Center Computational Cancer Biology Training Program of the Gulf Coast Consortia (funded by CPRIT Grant No. RP101489).

## Authors’ contributions

J.M. and Q.W. conceived and supervised the project. J.Z. and B.D.K. performed the prediction using *EpiPredictor*. J.D. performed all experimental characterizations. B.D.K. performed genome-wide analysis of K562 cell line. J.M. and Q.W. analyzed the results and wrote the paper. All authors agreed on the final manuscript.

